# Attention and Emotion-Enhanced Memory: A Systematic Review and Meta-Analysis of Behavioural and Neuroimaging Evidence

**DOI:** 10.1101/273920

**Authors:** Zachariah R. Cross, Amanda Santamaria, Mark J. Kohler

**Affiliations:** Centre for Cognitive and Systems Neuroscience, University of South Australia, Adelaide, Australia.; Sleep and Chronobiology Laboratory, University of South Australia, Adelaide, Australia.

**Keywords:** Attention, Emotion, Long-Term Memory, Electroencephalography, Functional Magnetic Resonance Imaging

## Abstract

The interaction between attention and emotion is posited to influence long-term memory consolidation. We systematically reviewed experiments investigating the influence of attention on emotional memory to determine: (i) the reported effect of attention on memory for emotional stimuli, and (ii) whether there is homogeneity between behavioural and neuroimaging based effects. Over half of the 47 included experiments found a moderate-to-large effect of attention on emotional memory as measured behaviourally. However, eye-tracking research provide mixed support for the role of attention-related processes in facilitating emotional information into long-term memory. Similarly, modulations in sensory-related components at encoding were not predictive of long-term memory formation, whereas later components appear to differentially reflect the allocation of attention to heterogeneous emotional stimuli. This dissociation in neurophysiology is paralleled by the activation of distinct neural networks under full- and divided-attention conditions. We quantified the effects of the behavioural, eye-tracking and neuroimaging findings via meta-analysis to show that the neural substrates of attention-related emotional memory enhancement may be sensitive to specific methodological parameters.

## 1. Introduction

The presence of an emotional element at encoding results in a memory trade-off, such that memory for emotional items are enhanced at the cost of memory for neutral information (Christianson, 1992; Riggs, McQuiggan, Farb, Anderson & Ryan, 2011). Early interpretations of emotion-enhanced memory are based on motivational models of emotion, which posit that the sensory system is constrained by a limitation in its processing capacity (Lang, Bradley & Cuthbert, 1998). Exogenous and endogenous attentional mechanisms allow the brain to deal with a subset of information that is deemed relevant or salient (LaBar & Cabeza, 2006; Okon-Singer, Hendler, Pessoa & Shackman, 2015; Vuilleumier, 2005). Thus, a trade-off in memory for emotional over neutral information may occur through attentional narrowing, whereby attention is rendered to emotional information for enhanced sensory processing (Chipchase & Chapman, 2013; Riggs et al., 2011). Mounting evidence supports this claim, suggesting emotional memory – the encoding and retrieval of an environmental or cognitive event that elicits an emotional response at the time of its occurrence – depends in part on attentional processes at encoding (Kensinger, 2009; Kensinger, Piguet, Krendl & Corkin, 2005; Riggs et al., 2011).

Behavioural studies illustrate that attention is modulated by emotion, whereby attention is drawn more rapidly to positive or aversive rather than neutral stimuli (Mackay et al., 2004; Kang, Wang, Surina & Lü, 2014). Studies demonstrating this finding often utilise the dot probe task (DPT), which requires subjects to simultaneously respond to stimuli varying in emotional valence (Mather & Carstensen, 2003). Research using this paradigm report slower reaction times for emotional compared to neutral stimuli of various modalities (e.g., auditory or visual stimuli) and of different stimulus types, such as images (Mather & Carstensen, 2003; Sakaki, Niki & Mather, 2012) and single words (Aquino & Arnell, 2007; Sharot & Phelps, 2004). Combined, this suggests emotional information is prioritised supramodally across sensory systems (Brosch & Van Bavel, 2012).

Electrophysiological (Carretié, Martín-Loeches, Hinojosa & Mercado, 2001; Schupp, Flaisch, Stockburger & Junghofer, 2006) and neuroanatomical studies (Sakaki et al., 2012; Smith, Henson, Rugg & Dolan, 2004; Smith, Stephan, Rugg & Dolan, 2006) report preferential neural activation patterns to emotional relative to neural stimuli from various brain regions. Electroencephalographic (EEG) research demonstrates a robust modulation in event-related potential (ERP) components by emotional stimuli (Zhang, Liu, An, Yang & Wang, 2015). Negative and positive compared to neutral stimuli elicit enhanced amplitudes in early sensory-related ERPs, and ERPs associated with elaborate stimulus evaluation, such as the P1 (Delplanque, Lavoie, Hot, Silvert & Sequeira, 2004), and late positive potential (LPP; Langeslag, Olivier, Kohlen, Nijs & Strien, 2015), respectively. The P1 is most pronounced at occipital regions and is modulated by attention (Delplanque et al. 2004), while the LPP is a positive-going waveform that is morphologically similar to the P3 (Luck, 2014). Analogous to the P3, the LPP may partially reflect the release of norepineprine from the brainstem Locus Coeruleus (LC), which innervates to the amygdala and hippocampal formation (Brown, Wee, Noorden, Giltay & Nieuwenhuis, 2015; Samuels & Szabadi, 2008). These innervations may facilitate cortical reorientation to emotionally significant events, promoting emotion-enhanced memory (Brown, Steenbergen, Band, Rover & Nieuwenhuis, 2012; Brown et al., 2015).

Neuroanatomical research (Keightley et al., 2003; Talmi, Anderson, Riggs, Caplan & Moscovitch, 2008; Vuilleumier et al., 2002) demonstrates that emotional memory is encoded and consolidated over neutral information due to interactions between the amygdala and fronto-parietal attentional networks, and the influence of the amygdala on the hippocampus during consolidation (Talmi et al., 2008; Taylor & Fragopanagos, 2005; Vuilleumier, 2005). In a meta-analysis of functional magnetic resonance (fMRI) studies (Murty, Ritchey, Adcock & LaBar, 2010), it was demonstrated that successful encoding of emotional information activated widespread neural networks, involving the ventral visual stream, hippocampal complex (i.e. anterior and posterior parahippocampal gyrus; PHG) and right ventral parietal cortex, as illustrated in Figure 1. These findings are analogous to magnetoencephalographic (MEG; Peyk, Schupp, Elbert & Junghofer, 2008; Keuper et al., 2014) research, which reveals emotional information is processed hierarchically, localised to occipital-parietal regions in early time windows (e.g., 120 – 170 ms), to more anterior, temporal regions in later time windows (e.g., 220 – 330 ms).

**Figure 1.**
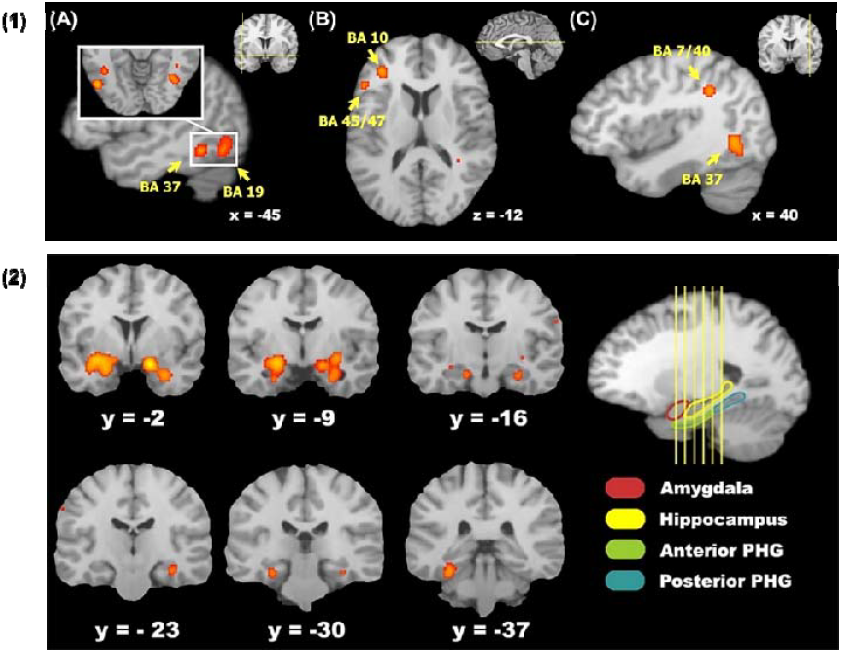
Activation likelihood map from a meta-analysis of 20 fMRI studies reported by Murty and colleagues (2010). **(1)** Regions showing reliable cortical activations in the ventral visual stream (A), the left prefrontal cortex; (B), and the right parietal cortex (C). **(2)** Regions showing reliable co-activation patterns between the amygdala and medial temporal lobe systems during encoding of emotional stimuli. Adapted from “fMRI studies of successful emotional memory encoding: A quantitative meta-analysis” by V. P. Murty, M. Ritchey, R. A. Adock, K. S. LaBar, 2010, *Neuropsychologia, 48*, p. 3462-3463, Copyright 2017 with permission from Elsevier (License number: 4232580453335).

There is compelling evidence to suggest attention and emotion interact to substantially influence sensory processing and memory consolidation (Schupp et al., 2006; Talmi et al., 2008). However, despite the breadth of research, behavioural and neuroimaging studies of the attention-emotional memory interface report mixed findings. Studies of eye movements demonstrate equal fixation time to negative and neutral items, while at recall, memory is enhanced for negative stimuli, but reduced for neutral items (Christiansan, Loftus, Hoffmann & Loftus, 1991; Riggs et al., 2011; Wessel, Van Der Kooy & Merckelbach, 2000). Further, electrophysiological findings, such as modulations in ERPs at encoding, often do not predict enhanced emotional memory performance at recall (Herbert et al., 2008; Keuper et al. 2014; Kissler, Herbert, Winkler & Junghofer, 2009).

To date there has been no attempt to systematically determine whether experimental research supports the role of attention in emotional LTM. In particular, a comprehensive review linking overt behavioural responses to changes in attention- and emotion-sensitive ERPs, and to the activation of cortical and subcortical brain networks has not yet been conducted. Systematically reviewing the behavioural and neuroimaging evidence of attention and emotional LTM may help to identify factors that characterise disorders that have an attentional preference for aversive events, such as anxiety and mood disorders (Lang et al., 2003; Leppänen, 2006). Such work would also help to illuminate the way in which the central nervous system prioritises information when its processing capacity is constrained, which may be particularly relevant for predictive-coding-based theories of brain function (e.g., Friston, 2010).

The current study aimed to determine whether attention significantly impacts memory for emotional stimuli. To this end, we systematically reviewed experiments investigating the influence of attention on emotional long-term memory (LTM) to determine: (i) if the literature demonstrates an effect of attention on memory for emotional stimuli; and (ii) whether there is homogeneity between behavioural, electrophysiological, and neuroanatomical correlates for the effect of attention on LTM for emotional information, and to quantify these effects via meta-analysis. A further aim of this review was to explore differences in the methodologies employed and provide suggestions for future emotional memory and attention research.

## 2. Method

This systematic review was conducted according to the Preferred Reporting Items for Systematic Reviews and Meta-Analyses (PRIMSA) guidelines (Liberati et al., 2009). PubMed, PsychInfo and Medline databases were searched on November 9, 2016. The search terms ‘Emotion*’, ‘Memory’, and ‘Attention*’ were used. A total of 8319 articles were identified and 2929 duplicates later removed. To be eligible for inclusion, articles had to be written in English, conducted with young adults (i.e., 18 – 35; Holland & Kensinger, 2012), included an assessment of emotional memory, and employed a measure of attention as it relates to memory processes (i.e., encoding, consolidation, or retrieval) for emotional stimuli. Further, only articles that were original, peer-reviewed research were included in the review. Articles from all publication years were accepted.

Exclusion criteria included studies with samples diagnosed with psychiatric disorders (e.g., depression, anxiety, and schizophrenia), studies conducted with infants and children (i.e., 17 years and under) or with middle and older adults (i.e., 36 years and above). However, where possible, data were extracted for the age group of interest from cross-sectional studies (e.g., Mather & Carstensen, 2003) that included a sample of young adults. Studies without full methodologies, without a measure of memory *and* attention, case studies and reviews, and/or studies that examined working memory as the measure of memory, were also excluded.

The primary reviewer (ZC) screened all titles and abstracts to determine eligibility. Articles that appeared to be eligible were sourced for full-text and reference lists of these articles were screened by title and abstract to ensure all eligible articles were included. Two reviewers (ZC and AS) screened eligible abstracts and full-text articles based on review criteria. Disagreements between reviewers were addressed through discussion; however, if the two reviewers did not reach a consensus, a third reviewer (MK) was consulted. The following data were extracted from all included articles: participant sample size, age and gender ratio, measure of attention (method and outcome measure), study characteristics (type of stimuli, study paradigm and recall interval) and major findings, separated by behavioural and neuroimaging (i.e., electrophysiological and neuroanatomical) results. Available effect sizes (i.e., *d*, □^2^, *r*, and *β*) were reported for relevant major findings.

### 2.1. Statistical analysis

Estimates of effect sizes were calculated using Pearson’s *r*, which is a valid effect size measure that is easily interpretable, and produces a measure between 0 (i.e., no relationship) and ±1 (perfect relationship; Field & Gillett, 2010). An *r* value of .10 is interpreted as a small effect, 0.30 as a moderate effect, and 0.50 as a large effect (Chatburn, Lushington, & Kohler, 2014; Cohen, 1992). The Hedges and Vevea (1998) random-effects model was used to calculate the meta-analysis. Random effects models are an appropriate method as they enable inferences to be made beyond the studies included in the meta-analysis, and are recommended to be the norm in psychological research (Field & Gillett, 2010). Of the 38 studies included in the systematic review, 32 provided data that were able to be converted into r values (i.e., *M* and *SD, F* statistic, *p* value, Cohen‘s *d*, □2, and β). In order to dissociate behavioural and neuroimaging effects, we calculated four separate random-effects models. In the first model, we included all 32 studies in order to obtain an overall effect of attention on emotional memory consolidation. We then calculated separate models for studies that employed behavioural, eye-tracking and neuroimaging (EEG/fMRI) measures. Analyses of publication bias were also computed, including Rosenthal’s fail safe *N*, and a random-effects publication bias model (Vevea & Woods, 2005).

## 3. Results

Figure 2 illustrates the flow of article selection, in accordance with Liberatie et al. (2009). 123 full-text articles were read for eligibility. 85 articles were excluded due to meeting one or more of the exclusion criteria. In total, 38 articles were included. A summary of the key study characteristics are provided in Table 1. Articles with behavioural and neuroimaging (i.e., EEG, fMRI) data are presented first, followed by articles consisting of single behavioural experiments. Articles with multiple eligible experiments are summarised last.

**Table 1.**
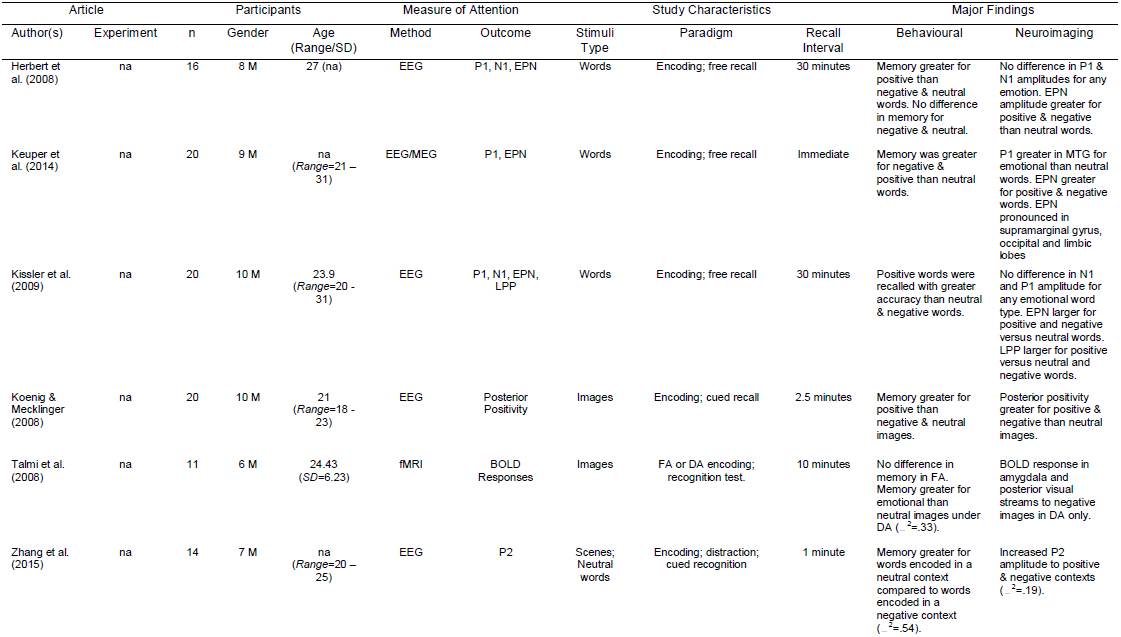
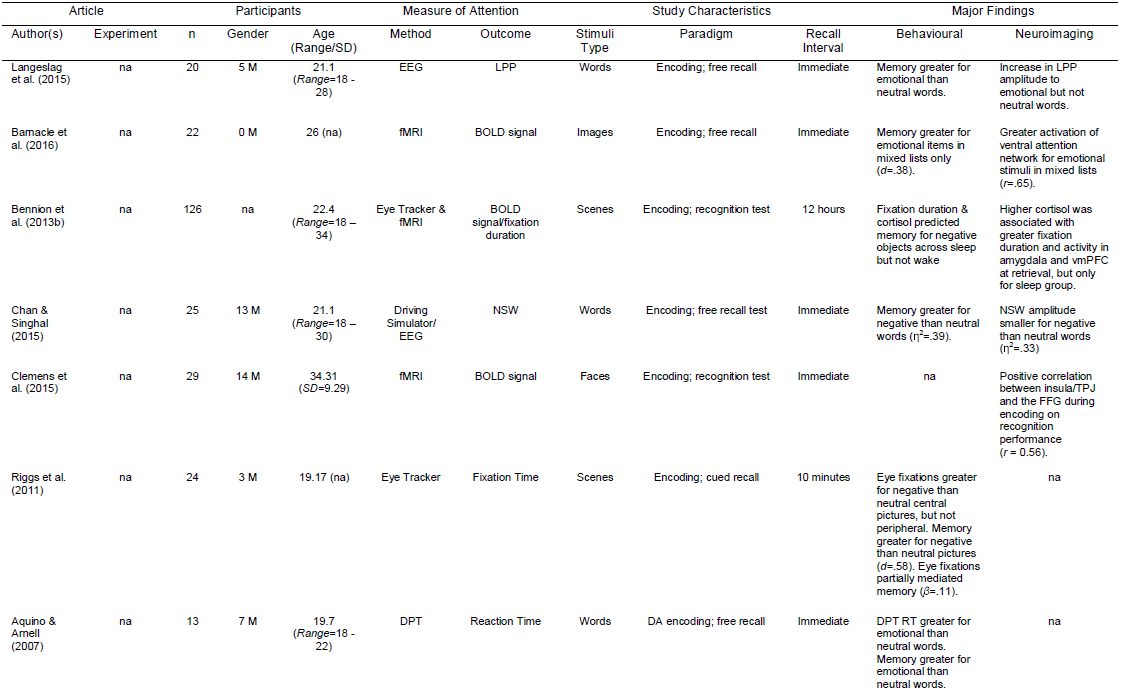
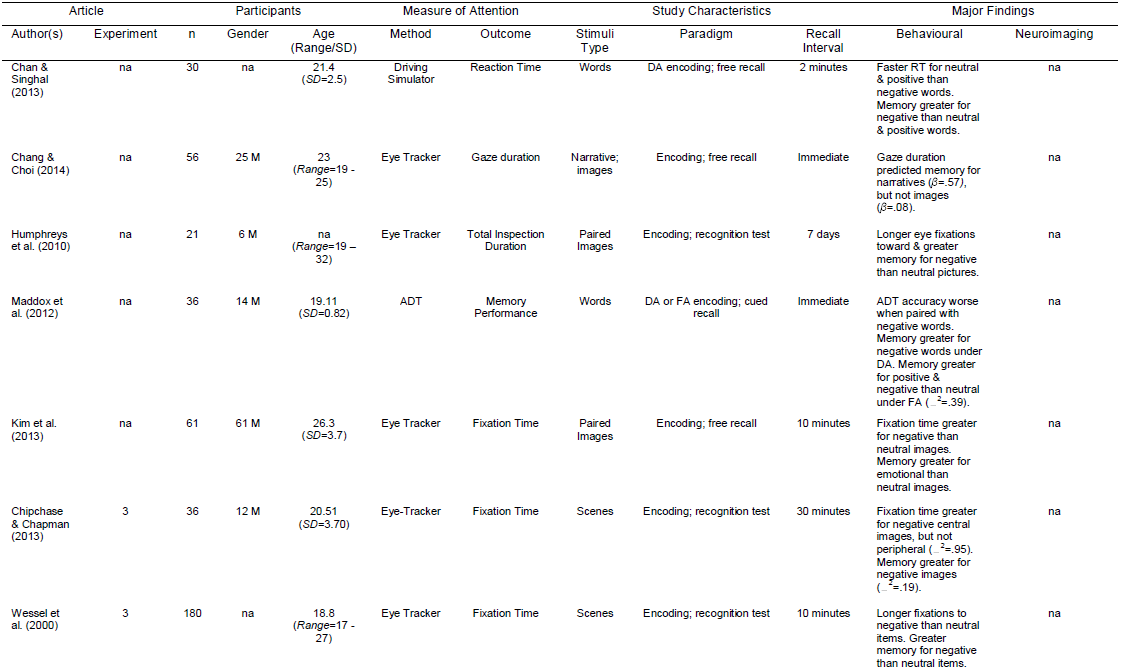
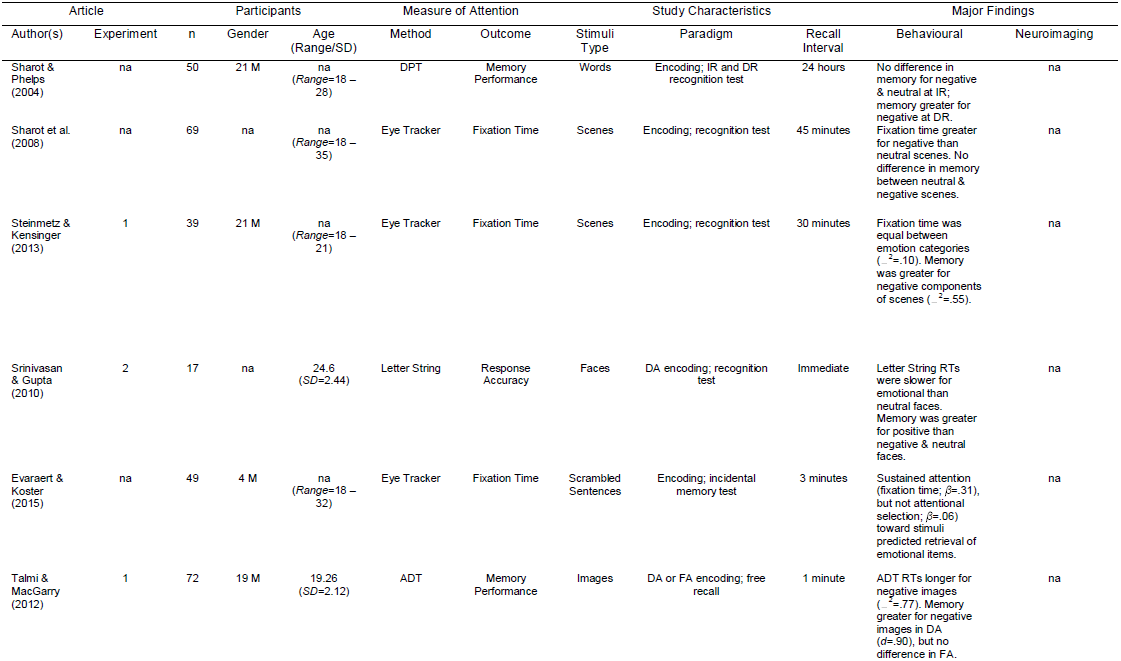
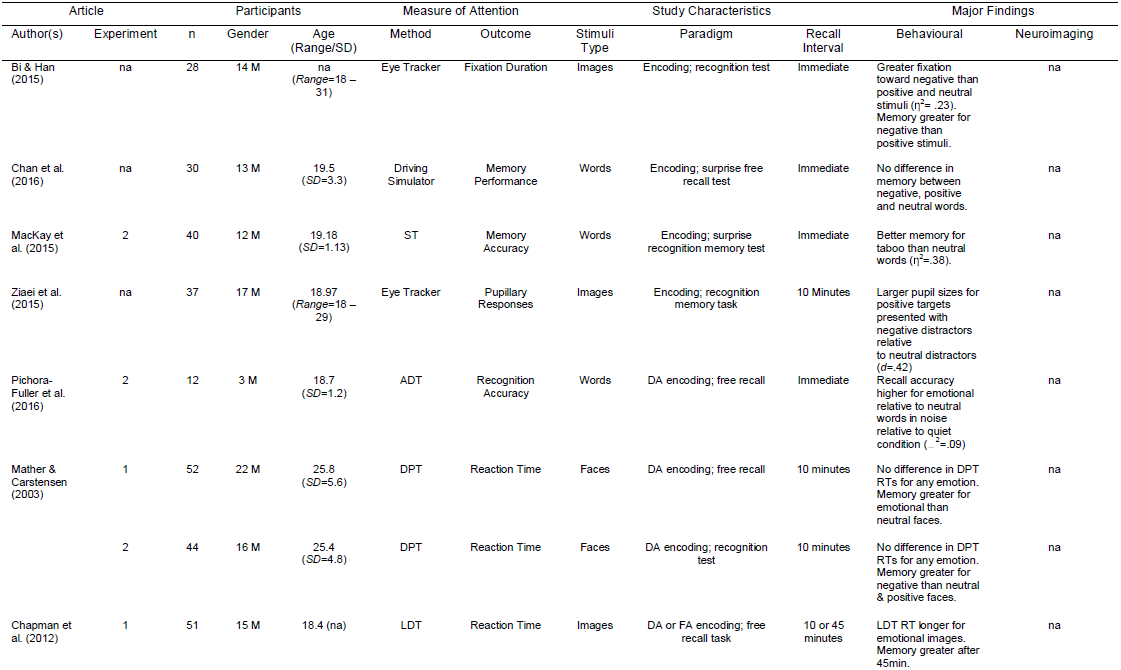
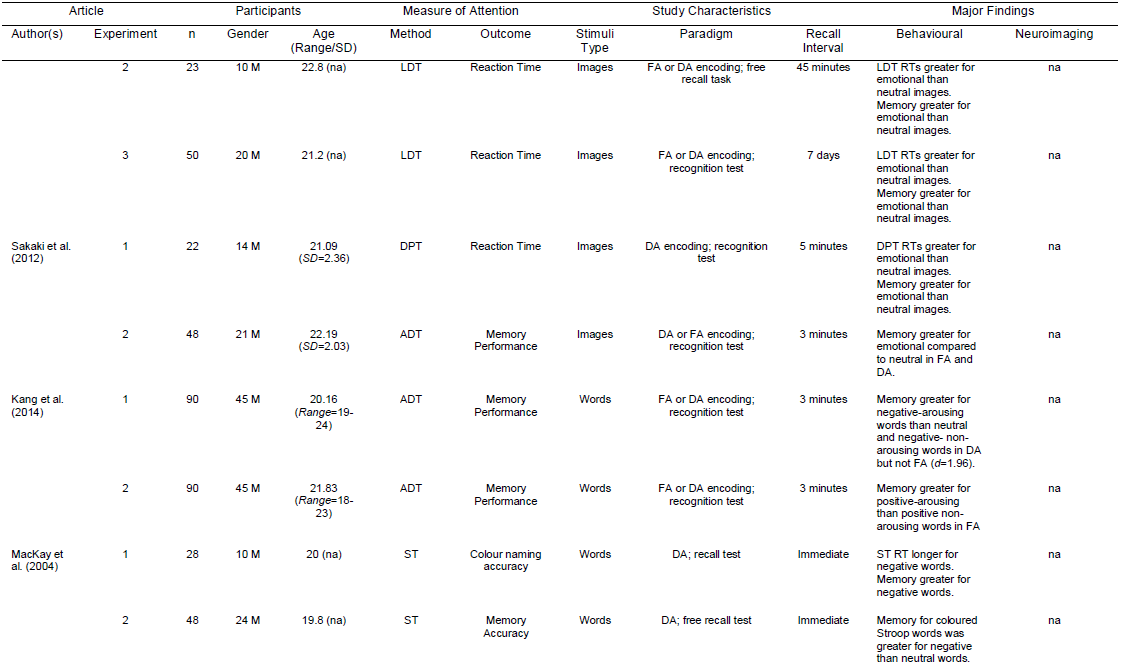
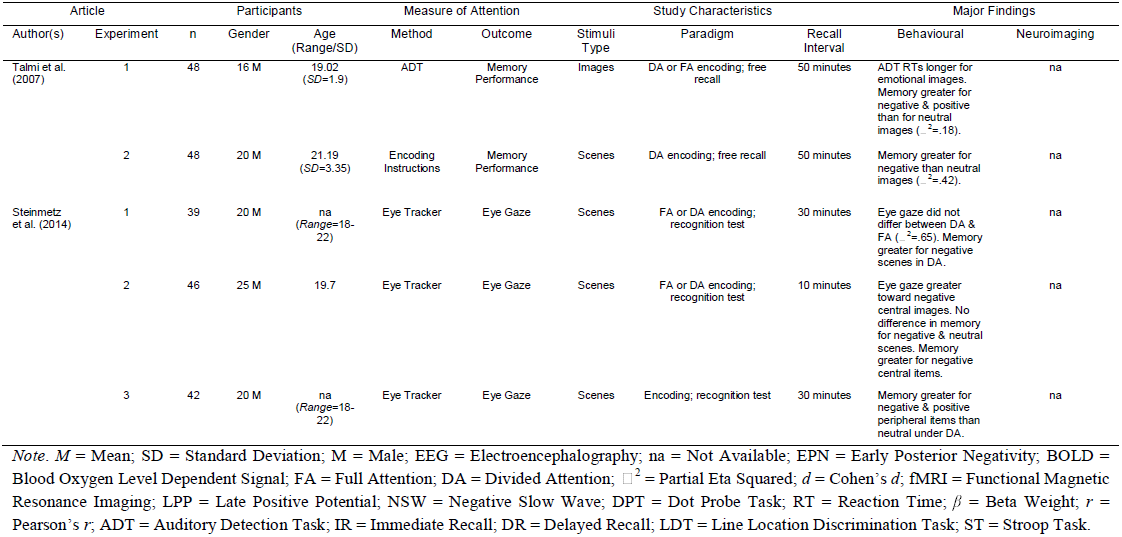
Summary of studies included in the systematic review ordered by neuroimaging, behavioural, and articles with multiple experiments.

**Figure 2.**
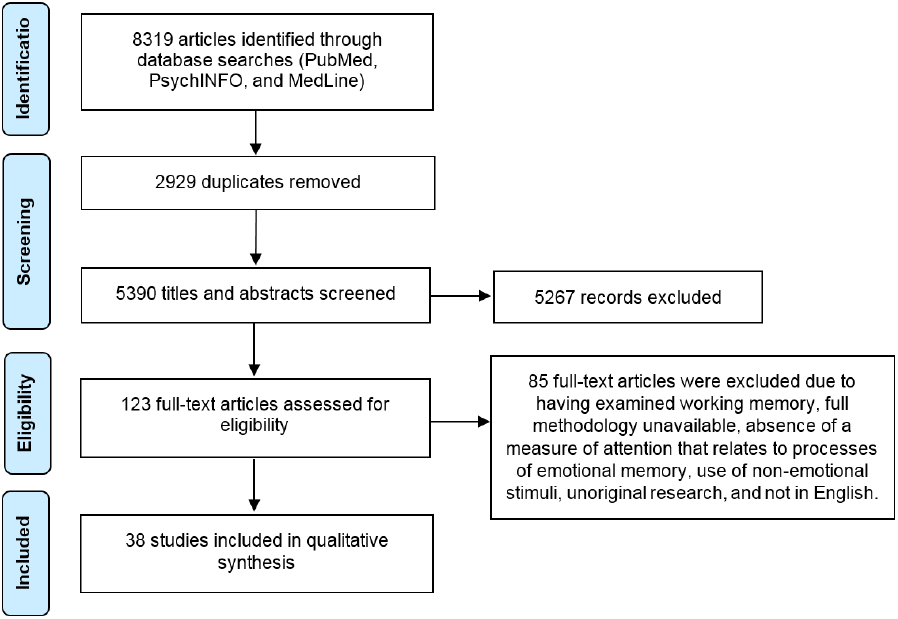
PRISMA flow diagram (Liberati et al., 2009) of the article screening and selection process. The databases searched included PubMed, PsychInfo, and MedLine.

Of the 38 articles reviewed, 7 reported multiple relevant experiments, resulting in 47 experiments being included. All experimental methodologies were examined and the following behavioural measures of attention were included: eye-tracking, DPT, driving simulator, ADT, encoding instructions provided by researchers, letter string, LDT, and the Stroop task.

This review identified 11 experiments that examined the neurobiological correlates of attention and emotional memory. Seven experiments used EEG to derive ERP components including the P1,N1, P2, early posterior negativity (EPN), posterior positivity and LPP, while one study used MEG (Keuper et al., 2014) to derive event-related magnetic fields (ERF). Four studies employed fMRI (e.g., Talmi et al., 2008) and measured the BOLD signal to index attention to emotional stimuli at encoding.

### 3.1. Measures of emotional memory

One study used stimuli from the Affective Norms for English Words (ANEW; e.g., Chan & Singhal, 2013), while twelve studies used the International Affective Picture System (IAPS; e.g.,Chapman et al., 2012; Humphreys et al., 2010; Koenig & Mecklinger, 2008; Riggs et al., 2011; Talmi & MacGarry, 2012; Talmi et al., 2007; Talmi et al., 2008; Zhang et al., 2015). Three studies used emotionally valenced faces (e.g., Clemens et al., 2015; Mather & Carstensen, 2003; Srinivasan & Gupta, 2010), while one study used images and emotional narratives (Chang & Choi, 2014). Remaining studies (30) derived stimuli from previous studies (e.g., established sets used by Payne, Stickgold, Swanberg & Kensinger, 2008) or Google Images. All studies without standardised stimuli generated inter-rater reliability estimates from expert judges and/or gained valence and arousal ratings from participants (e.g., Maddox et al., 2012; Sakaki et al., 2012).

Two studies (Humphreys et al., 2010; Kim et al., 2013) presented pairs of images (e.g., neutral-negative, neutral-neutral, positive-negative, and positive-neutral) to participants at encoding. Twelve experiments used emotional scenes, involving a central negative or neutral item with a neutral or negative peripheral background image, respectively. The remaining 33 experiments presented single stimuli for participants to learn.

### 3.2. Paradigms

Twenty-four experiments used a passive encoding paradigm, requiring participants to view stimuli passively for a set time with stimulus presentation ranging from one to six seconds. Of the 47 experiments reviewed, 23 had full- and divided-attention paradigms, requiring participants to encode stimuli while engaging in a second task (e.g., DPT or ADT). Twenty-six experiments used cued recognition tests to assess participants’ memory, involving participants responding to previously seen stimuli intermixed with new distractor stimuli. Twenty-one of the 47 experiments used a free recall paradigm, requiring participants to write down as many stimuli as they could remember with no time constraints.

### 3.3. Recall intervals

Thirty-two percent of studies tested immediate recall performance, 19% had a recall interval of less than five minutes, 32% of studies had a recall interval of between ten and thirty minutes, while 8.5% of studies employed a recall interval of between 45 – 60 minutes and 8.5% of greater than 60 minutes. All studies using short recall interval (i.e., <5 minutes – 60 minutes) had participants complete filler tasks, such as simple arithmetic.

### 3.4. Behavioural measures of attention

#### 3.4.1. Auditory discrimination task

Seven experiments used an ADT to assess the effect of DA at encoding of emotional memory. All experiments reported significant effects of attention, such that when attention was divided between encoding and the ADT, memory was greater for negative than neutral and positive stimuli. Three experiments (Maddox et al., 2012; Kang et al., 2014; Talmi & MacGarry, 2012) reported greater memory for negative than neutral and positive stimuli under DA, but greater memory for negative and positive than neutral stimuli under full attention (FA).

#### 3.4.2. Dot probe task

Five experiments examined the effect of the DPT on emotional memory. Two studies (Aquino & Arnell, 2007; Sakaki et al., 2012) reported longer reaction times to negative than neutral stimuli, which translated into greater memory for negative stimuli at immediate recall and after a five minute recall interval. One study (Sharot & Phelps, 2004) reported no difference in memory for negative and neutral words at immediate recall; however, when memory was tested after 24 hours, memory was enhanced for negative stimuli, but diminished for neutral stimuli.

One study containing two experiments (Mather & Carstensen, 2003) reported no statistically significant difference in DPT reaction times for negative and neutral faces. However, memory was significantly greater for negative faces after a ten minute recall interval. This finding was replicated using positive faces, whereby memory for negative faces was greater than memory for neutral and positive faces.

#### 3.4.3. Driving simulator

Three studies used a driving simulator to test attention to emotional items (Chan & Singhal, 2013, 2015; Chan et al., 2016). Participants were required to respond via a button on a steering wheel when a target stimulus (neutral word) appeared on a billboard. Reaction times were slower when the target stimulus was paired with negative and positive words. In two of the studies (Chan & Singhal, 2013, 2015), memory was greater for positive and negative than neutral words after a two minute recall interval. The study with non-significant results (Chan et al., 2016) tested participants’ memory immediately after the encoding phase.

#### 3.4.4. Encoding instructions

Using emotional scenes, one experiment (Talmi et al., 2007) instructed participants to pay attention to central items and to ignore peripheral objects. However, memory was significantly greater for central and peripheral negative objects than central and peripheral neutral objects after a 50 minute recall interval.

#### 3.4.5. Eye-tracking

Fifteen experiments used eye-tracking as a measure of overt attention. Two experiments (Humpreys et al., 2010; Kim et al., 2013) using paired-pictures reported greater fixations to negative than neutral images, which translated into enhanced memory for negative over neutral items at recall. One experiment (Chang & Choi, 2014) reported that gaze duration predicted memory for emotional narratives but not for emotional images, while another study (Bi & Han, 2015) found that greater fixation time toward negative than positive and neutral stimuli predicted memory performance for negative stimuli. Of the 15 experiments, one (Ziaei et al., 2015) measured pupillary responses, and reported larger pupil sizes for positive targets presented with negative distractors relative to neutral distractors, which translated into greater memory for positive items.

Of the seven studies using emotional scenes, one experiment (Sharot et al., 2008) reported no difference in memory between negative and neutral stimuli, despite greater fixations for negative than neutral central and peripheral objects at encoding. The remaining six studies (Bennion et al., 2013b; Chipchase & Chapman, 2013; Riggs et al., 2011; Steinmetz & Kensinger, 2013; Steinmetz et al., 2014; Wessel et al., 2000) reported longer fixations to negative than neutral central items, but no difference in the length of fixations to negative and neutral peripheral images. All six studies reported greater memory performance for negative, relative to neutral, central and peripheral items. However, one study (Bennion et al., 2013b) also included a sleep group (i.e. compared sleep versus wake interval between learning and retrieval), and found that fixation duration predicted memory for negative objects across sleep but not wake.

In two of three experiments reported by Steinmetz et al. (2014), gaze duration did not differ between negative, neutral and positive scenes in the DA condition. After a 30 minute recall interval, memory performance was greater for negative compared to positive, neutral peripheral and central items. In the third experiment, gaze duration was greater toward negative central, but not peripheral items; however, after a ten minute recall interval, memory performance was greater for central and peripheral negative items than central and peripheral neutral items.

The one study (Evaraert & Koster, 2015) that examined the effect of attention (i.e., fixation time) on retrieval of emotional memory reported a significant effect of sustained attention toward positive and negative items, but a non-significant effect of attentional selection on recollection of emotional words embedded in scrambled sentences.

#### 3.4.6. Letter string

One study used the Letter String task to assess the effect of attention on memory for emotional faces (Srinivasan & Gupta, 2010). Participants were required to indicate on a button box whenever the letter ‘N’ was present in a string of letters that were superimposed on a face. Letter string reaction times were significantly longer for negative and positive compared to neutral faces; however, memory was greater for positive than neutral and negative faces at immediate recall.

#### 3.4.7. Line location discrimination task

One article (Chapman et al., 2012) reporting three experiments used the line LDT; a computerised task requiring participants to indicate whether a line appears above or below a stimulus. In all three experiments, negative images elicited slower LDT reaction times relative to neutral stimuli. Further, memory was greater for negative than neutral stimuli at all three recall intervals (10 minutes, 45 minutes, and 7 days). However, memory for negative images was equivalent after a 45 minute delay relative to 10 minute recall interval and this effect maintained after a seven day recall interval. Memory for neutral items was reduced after the 45 minute and 7 day recall intervals relative to the 10 minute recall interval.

#### 3.4.8. Stroop task

Two articles used the Stroop task to assess the role of attention in emotional memory (MacKay et al., 2004, 2015). In both studies, participants were required to name the colour of neutral and negative words. In the first study (MacKay et al., 2004), reaction times were longer for negative than neutral words at encoding, which led to enhanced memory for negative words at immediate recall. In the second study (MacKay et al., 2015), memory was better for negative versus neutral words.

### 3.5. Electrophysiological and neuroanatomical measures of attention

#### 3.5.1. Event-related potentials/fields

Three studies investigated modulations in P1 in response to emotional words (Herbert et al., 2008; Keuper et al., 2014; Kissler et al., 2009). No study reported statistically significant differences in the amplitude of the P1 between negative, neutral, and positive word types. All three experiments reported greater memory for positive than neutral, and negative words after a 30 minute (Herbert et al., 2008; Kissler et al., 2009) and immediate (Keuper et al., 2014) recall interval. Two of these three experiments examined modulations in N1 amplitude to emotional words (Herbet et al., 2008; Kissler et al., 2009). Both studies reported no statistically significant difference in N1 amplitude to positive, negative, and neutral words.

One study reported significant effects for the P2, such that positive and negative images elicited an increase in P2 amplitude relative to neutral images (Zhang et al., 2015). Memory for neutral words that were superimposed on positive and negative images was poorer compared to neutral words superimposed on neutral images, demonstrating an emotion-enhanced memory trade-off.

Two studies examined the EPN (Herbert et al., 2008; Keuper et al., 2014). Both studies employed emotional words and reported greater EPN amplitude toward positive and negative than neutral words. There was no difference in EPN amplitude between positive and negative words. Using MEG, Keuper et al. (2014) reported greater ERFs within the time window of the P1 (80 – 120ms) in the middle temporal gyrus for emotional compared to neutral words. Greater ERFs within the EPN time window (200 – 300ms) were observed in the supramarginal gyrus, occipital lobe (cuneus and precuneus) and limbic lobe (posterior cingulate cortex) for emotional compared to neutral words.

One study reported increased posterior positivity amplitude to positive and negative versus neutral images, which correlated with greater memory performance for positive and negative images (Koenig & Mecklinger, 2008). Similarly, two experiments examined the LPP and reported increases in LPP amplitude to emotional relative to neutral words (Kissler et al., 2009; Langeslag et al., 2015). Memory performance was greater for emotional than neutral words at immediate recall. Finally, one experiment (Chan & Singhal, 2015) reported reduce negative slow wave (NSW) amplitude for negative versus neutral words, which was coupled with enhanced memory for negative over neutral words at immediate recall.

#### 3.5.2. Functional magnetic resonance imaging

Using fMRI, four studies measured the blood oxygen-level dependent (BOLD) signal to index attention to emotional stimuli. Talmi et al. (2008) reported greater BOLD responses in the amygdala and posterior visual streams to negative than for neutral images under DA but not under FA. There was no difference in memory between negative and neutral images under FA, but greater memory for emotional than neutral images under DA was reported. Similarly, Barnacle et al. (2016) reported greater activation of the ventral attention network during the encoding of emotional images in mixed (i.e. emotional mixed with neutral stimuli) but not pure lists, which was associated with enhanced memory for emotional over neutral items at recall.

Clemens et al. (2015) reported positive correlations between activity in the insula/TPJ and FFG during encoding on recognition memory. Finally, using concurrent eye-tracking, cortisol assaying and fMRI, Bennion et al. (2013b) found that higher cortisol during encoding was associated with greater fixation duration toward negative stimuli, which predicted activity in the amygdala and vmPFC at retrieval. Interestingly, this effect was only present for across a period containing sleep, such that memory for negative items was greater after a period of sleep versus wake, and this was mediated by cortisol levels at encoding.

### 3.6. Meta-analysis

The meta-analysis contained 32 studies, comprising an overall sample of 1291 young adults. As per Brewin et al. (2007), we averaged effects for studies that reported multiple relevant effect sizes, so that each experiment contributed a single data point to the overall model (see Table 3 for a summary of the studies included in the meta-analysis). The mean pooled effect size for attention on emotional long-term memory was .40 (*p* <.001, 95% CI = [.32, .48]). This indicates a moderate effect, with a significant z of 9.058, *p*<.001. Separate analyses revealed a moderate-to-large mean effect for behavioural (*r* = .46, *p* <.001, 95% CI = [.29, .54], z = 5.704, *p*<.001) and neuroimaging (*r* = .54, *p* <.001, 95% CI = [.40, .65], z = 6.777, *p*<.001) studies, and a small-to-moderate mean effect for eye-tracking studies (*r* = .31, *p* <.001, 95% CI = [.19, .42], z = 5.003, *p*<.001). Note that chi-square tests for homogeneity of variance were not significant for any of the models (see Table 2 for a summary of the homogeneity tests for each model). However, based on recommendations by Field and Gillett (2010) and Chatburn et al. (2014), random-effects models were used, as there is likely to be large variance in attention for various types of emotional stimuli – both behaviourally and neurobiologically – in the larger population. Thus, the insignificant chi-square tests are likely to reflect the small number of studies (32), rather than a lack of variability between experiments. Meta-analysis results are illustrated in Figure 4.

**Table 2.** Chi-square tests for homogeneity of variance for the four random-effects models.

| Model | $\chi^2$ | $df$ | $p$ | $n$ |
| --- | --- | --- | --- | --- |
| Main | 28.37 | 31 | 0.602 | 32 |
| Behavioural | 10.26 | 12 | 0.592 | 13 |
| Eye-Tracking | 10.76 | 11 | 0.463 | 12 |
| Neuroimaging | 4.07 | 6 | 0.667 | 7 |

Note. $n$ = number of studies.

**Table 3.** Sample size, effect size, lower and upper 95% confidence intervals, measure, stimuli and recall interval for studies included in the meta-analysis. Studies are ranked based on largest to smallest effect size.

| Study | *n* | *r* | Lower 95% CI | Upper 95% CI | Measure | Stimuli | Recall Interval |
| --- | --- | --- | --- | --- | --- | --- | --- |
| Talmi et al. (2008) | 11 | 0.80 | 0.38 | 0.94 | fMRI | Images | 10 minutes |
| Talmi et al. (2007) | 48 | 0.73 | 0.56 | 0.84 | Instructions | Images | 50 minutes |
| Kang et al. (2014) | 90 | 0.69 | 0.56 | 0.78 | ADT | Words | 3 minutes |
| Barnacle et al. (2016) | 22 | 0.65 | 0.31 | 0.84 | fMRI | Images | Immediate |
| Chang & Choi (2014) | 56 | 0.64 | 0.45 | 0.77 | Eye-Tracking | Words | Immediate |
| Chan & Singhal (2015) | 25 | 0.57 | 0.22 | 0.78 | EEG | Words | Immediate |
| Clemens et al. (2015) | 29 | 0.56 | 0.24 | 0.76 | fMRI | Images | Immediate |
| Chipchase & Chapman (2013) | 36 | 0.53 | 0.25 | 0.73 | Eye-Tracking | Images | 30 minutes |
| Srinivasan & Gupta (2010) | 17 | 0.48 | 0.00 | 0.78 | Letter String | Images | Immediate |
| MacKay et al. (2004) | 28 | 0.47 | 0.12 | 0.72 | ST | Words | Immediate |
| Bi & Han (2015) | 28 | 0.47 | 0.12 | 0.72 | Eye-Tracking | Images | Immediate |
| MacKay et al. (2015) | 40 | 0.44 | 0.24 | 0.61 | ST | Words | Immediate |
| Koenig & Mecklinger (2008) | 20 | 0.44 | 0.00 | 0.74 | EEG | Images | 2.5 minutes |
| Zhang et al. (2015) | 14 | 0.43 | -0.13 | 0.78 | EEG | Words | 1 minute |
| Talmi & MacGarry (2012) | 72 | 0.41 | 0.19 | 0.58 | ADT | Images | 1 minute |
| Sakaki et al. (2012) | 48 | 0.41 | 0.14 | 0.62 | ADT | Images | 3 minutes |
| Steinmetz et al. (2014) | 39 | 0.41 | 0.11 | 0.64 | Eye-Tracking | Images | 30 minutes |
| Sharot & Phelps (2004) | 25 | 0.40 | 0.01 | 0.68 | DPT | Words | 24 hours |
| Chan & Singhal (2013) | 30 | 0.39 | 0.15 | 0.59 | EEG | Words | Immediate |
| Bennion et al. (2013b) | 24 | 0.37 | -0.03 | 0.67 | Eye-Tracker/fMRI | Images | 12 hours |
| Sharot et al. (2008) | 69 | 0.33 | 0.10 | 0.52 | Eye-Tracking | Images | 45 minutes |
| Steinmetz & Kensinger (2013) | 36 | 0.31 | -0.01 | 0.58 | Eye-Tracking | Images | 30 minutes |
| Pichora-Fuller et al. (2016) | 12 | 0.30 | -0.33 | 0.74 | ADT | Words | Immediate |
| Maddox et al. (2012) | 36 | 0.28 | 0.05 | 0.55 | ADT | Words | Immediate |
| Kim et al. (2013) | 61 | 0.23 | -0.02 | 0.45 | Eye-Tracking | Images | 10 minutes |
| Evaraert & Koster (2015) | 49 | 0.20 | -0.08 | 0.45 | Eye-Tracking | Words | 3 minutes |
| Ziaei et al. (2015) | 37 | 0.20 | -0.13 | 0.49 | Eye-Tracking | Images | 10 minutes |
| Mather & Carstensen (2003) | 44 | 0.13 | -0.07 | 0.32 | DPT | Images | 10 minutes |
| Riggs et al. (2011) | 24 | 0.10 | -0.35 | 0.52 | Eye-Tracking | Images | 10 minutes |
| Wessel et al. (2000) | 180 | 0.05 | -0.23 | 0.34 | Eye-Tracking | Images | 10 minutes |
| Humphreys et al. (2010) | 18 | -0.01 | -0.47 | 0.45 | Eye-Tracking | Images | 7 days |
| Chapman et al. (2012) | 23 | -0.06 | -0.46 | 0.35 | LDT | Images | 45 minutes |
| **Meta-Analytic Average for Main Effect** | **1291** | **.40** | **[-0.32, 0.48]** |  |  |  |  |

*Note.* *n* = sample size; *r* = correlation coefficient; CI = confidence interval; fMRI = functional magnetic resonance imaging; EEG = electroencephalography; ADT = auditory detection task; ST = Stroop task; DPT = dot probe task; LDT = line dissection task. The dashed lines separate studies with high (>0.50), moderate (0.30 – 0.49) and low (<0.29) effect sizes.

**Figure 3.**
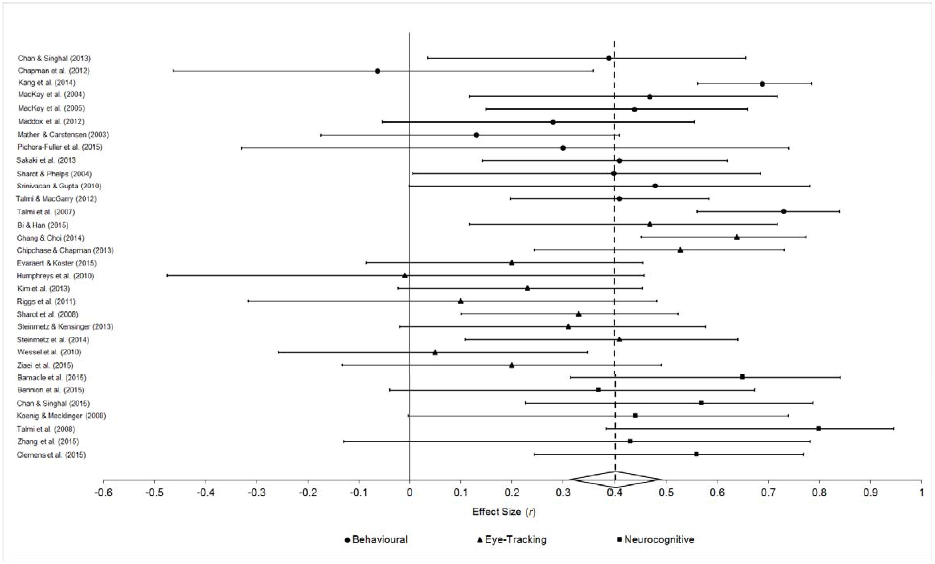
Forest plot for the effect (*r*) of attention on emotional long-term memory for behavioural, eye-tracking and neuroimaging studies. Circles indicate behavioural findings: triangles indicate eye-tracking findings; squares indicate neuroimaging (EEG/fMRI) findings. Bars indicate the 95% confidence intervals of each effect. The studies corresponding to each effect are listed on the left. The diamond and dashed line located at the bottom of the figure indicate the mean meta-analytic effect size with 95% confidence intervals.

### Publication bias

Several analyses were performed to examine potential publication bias. Rosenthal’s fail safe *N* indicated that 1936 studies with negative findings would need to be excluded for the population effect size estimate to be non-significant. To further examine potential publication bias, a random-effects publication bias model (Vevea & Woods, 2005) was computed. A moderate two-tailed selection bias model reduced the overall effect size to 0.37, while a severe two-tailed selection bias model yielded an adjusted effect size of 0.34. From this perspective, if there were publication biases in the meta-analytic sample, the effects would be minimal, given that the adjusted effect size estimates remained moderate.

## 4. Discussion

A total of 47 experiments across 38 studies were included in this systematic review, 32 of which were subject to meta-analysis. These studies utilised various behavioural, electrophysiological, and neuroanatomical measures to investigate the effect of attention on emotional memory consolidation. Of the behavioural studies included, over half reported statistically significant effects of attention on emotional memory, suggesting attention modulates emotional memory consolidation. The use of DA paradigms (e.g., Maddox et al., 2012; Kang et al., 2014; Talmi & MacGarry, 2012) provide the majority of behavioural evidence for these effects. However, we found largely inconsistent evidence for the role of overt attention in emotional memory for experiments utilising eye-tracking, which were corroborated by ERP experiments that reported null effects in early sensory-related potentials, but modulations in later components. A discussion of the discrepancy between the behavioural, electrophysiological and neuroanatomical findings is presented below, followed by suggestions for future research.

### 4.1. Behavioural effects of attention on emotional memory

The greatest effects on emotional memory were observed in DA paradigms, where participants attended to a secondary task during encoding. Experiments utilising the ADT (e.g., Maddox et al., 2012) and DPT (e.g., Aquino & Arnell, 2007) reported longer reaction times in response to emotional relative to neutral stimuli, suggesting attentional resources are preferentially allocated to salient relative to neutral information for elaborate evaluation. Across experiments, longer reaction times resulted in enhanced memory performance at recall. This emotional slowing may be explained by inhibitory effects of emotion on selective attention, whereby attention is inhibited by emotional cues, resulting in a slowing of responses to secondary tasks, such as the ADT and DPT (Bradley, 2009). This provides support for an attentional narrowing account of emotional memory, wherein emotional information interferes with cognitive and motor goals that prepare responses to incoming stimuli, reducing evaluation of less salient information and facilitating emotional information into LTM (Taylor & Fragopanagos, 2005).

Studies measuring eye movements suggest attention does not entirely account for the effect of emotion on memory. Utilising fixation time as a measure of overt attention, two experiments conducted mediation analyses to test the relationship between attention and emotional memory (Kim et al., 2013; Riggs et al., 2011). Both studies reported that the direct path between emotion and memory for central items remained significant when attention was fixed. Further analyses revealed that the indirect path between attention and memory for peripheral objects was not significant, such as that memory for negative peripheral objects remained greater than for neutral peripheral objects, irrespective of processing time (e.g., fixation time).

This discrepancy between eye movements and behavioural responses lends support to the argument that post-encoding processes are more critical for the consolidation of emotional information than heightened attention at encoding (Kim et al., 2013; Riggs et al., 2011). However, it is potentially problematic to assume that eye movements can be used as direct measures of overt attention, and thus predictors of behavioural outcomes. From a predictive-coding-based view of the brain, eye movements serve to gather data from the environment to test beliefs about the current internal model of the world (Friston et al., 2012). This account of eye movements is based on the free-energy principle, which posits biological agents act upon the world according to encoded representations of sensations, and that adaptive responses to the environment occur under conditions within which sampled sensations match internal predictions (Friston, 2010; Joffily & Coricelli, 2013). An emotional stimulus that is congruent with an individual’s internal model may therefore elicit shorter fixations than a stimulus that is not, suggesting emotional information may serve to regulate the learning rate of a biological agent rather than directly reflecting modulations in attentional orientation (Joffily & Coricelli, 2013). Therefore, although attentional narrowing models are consistent with behavioural effects, more research is needed to understand the brain mechanisms underpinning the role of attention in emotional LTM.

### 4.2. Neural basis of attention-related emotional memory enhancement

Although there is a breadth of research reporting the behavioural correlates of attention and emotional memory, few experiments have investigated the electrophysiological and neuroanatomical correlates of this relationship. Eleven (23%) experiments included in this review examined the electrophysiological and neuroanatomical correlates of attention and emotional memory. Modulations in sensory-related ERP components at encoding were not related to behavioural performance at recall. All studies examining modulations of the P1 and N1 in response to emotional words at encoding reported nonsignificant effects of emotion on their amplitude or latency (Herbert et al., 2008; Kissler et al., 2009). Prior research demonstrates that the P1 and N1 are modulated when attentional state is under demand (for review: Schupp et al., 2006), suggesting single word-based stimuli and scene imagery may be better indexed by later components, such as the P2 and EPN (Herbert et al., 2008).

The P2 is suggested to index post-perceptual selective attention and is sensitive to visual target detection, such that P2 amplitude has been reported to increase when target stimuli are detected among distractor objects (Kanske, Plitschka & Kotz, 2011). In one experiment, P2 amplitude increased in response to emotional words imbedded in complex visual scenes, and was associated with superior memory performance for emotional relative to neutral stimuli at recall (Zhang et al., 2015). In this experiment, neutral words were superimposed on emotionally valenced backgrounds, a paradigm more complex than the passive paradigms utilised by studies investigating the P1 and N1. The P2 may therefore represent a top-down attentional influence on emotion evaluation, particularly when presented alongside neutral stimuli, and may index attention-related cortical networks that facilitate emotional information into LTM (Carretié et al., 2004; Crowley & Colrain, 2004).

The EPN proceeds the P2 and is thought to index selective attention to specific stimulus features (Junghöfer, Bradley, Elbert & Lang, 2001). Included studies reported enhanced EPN amplitude to emotional relative to neutral stimuli, suggesting increased visual processing of emotional stimuli, and a prioritisation of emotional over neutral information. However, these components may reflect post-attentive, elaborate stimulus evaluation, rather than early attentional orientation (Low, Lang & Bradley, 2005; Thorpe, Fize & Marlot, 1996). For example, Low et al. (2005) systematically demonstrated that the EPN is modulated by picture content (objects vs people) and picture type (central vs peripheral items) above that of emotionality, suggesting the EPN reflects selective attention rather than the automatic detection and evaluation of emotion. However, the studies examining the EPN included in the meta-analysis used emotionally valenced words, and each reported greater EPN amplitude in response to emotional compared to neutral words. Moreover, using MEG, Keuper et al. (2014) reported that the EPN was generated from occipital and limbic lobes, suggesting possible functional connectivity between primary visual cortices and the amygdala. This may subserve selective attention to, and the preferential processing of emotional information, which accords with fMRI research (for review: Murty et al., 2010).

In contrast to sensory-related components (i.e. N1, P1), the amplitudes of the posterior positivity and LPP were larger in response to emotional relative to neutral stimuli. The posterior positivity has been hypothesised to reflect attentional capturing mechanisms during early stages of the processing cascade (i.e. 250 – 450 ms; Koenig & Mecklinger, 2008). Similarly, the LPP has been likened to a P3b response, in the sense that the LPP is considered a sensitive measure of attentional orientation to salient information (Schupp et al., 2006). Importantly, the studies investigating the modulation of sensory-related components did not manipulate attention, and therefore, under a biased competition model of attention, there was no demand on processing resources in the relevant sensory cortex (i.e. visual or auditory cortices). This is in contrast to the posterior positivity and LPP, where irrespective of attentional conditions, they were still modulated by emotion. From this perspective, the N1 and P1 may serve to gate incoming sensory information via bottom-up processing mechanisms. During later processing stages, components such as the LPP may facilitate enhanced memory via the release of norepineprine from the LC in response to emotionally salient stimuli, which would promote synaptic plasticity via afferent inputs to the hippocampal complex and amygdala (Tully & Bolshakov, 2010).

All fMRI studies reported significant effects (*r* = 0.37 – 0.80) of attention on emotional memory, and revealed a distributed network of cortical and subcortical regions during the encoding of emotional stimuli under varying attentional conditions. Studies consistently revealed greater bilateral amygdala activation in response to emotional relative to neutral stimuli. When attention was divided, two studies (Barnacle et al., 2016; Tamli et al., 2008) reported greater activation of the ventral attention network, which predicted enhanced memory performance at recall for emotional relative to neutral stimuli. The ventral attention network – which includes the anterior insula, anterior cingulate cortex and temporo-parietal junction – is involved in reorienting attention to task-relevant stimuli (Corbetta & Shulman, 2002; Fragopanagos & Taylor, 2006). However, greater activation of the ventral attention network in response to emotional stimuli under DA accords with evidence indicating that this processing stream is susceptible to emotionally salient information irrespective of task-relevance (for review: Frank & Sabatinelli, 2012). From this perspective, during top-down, goal-directed processing, the ventral attention network is critical for detecting bottom-up, stimulus-driven changes; however, emotional stimuli appear to have privilege access to attention, which in turn, promotes emotion-enhanced memory. This interpretation is consistent with behavioural experiments that report memory for emotional stimuli is enhanced under DA compared to FA, and lends support to the argument that attention is a strong predictor for long-term emotional memory consolidation.

Taken together, attention at encoding is a strong predictor of long-term emotional memory. Effects of attention are moderate at the behavioural level, while the largest effects are born from neuroimaging (EEG/fMRI) studies. In contrast, while still significant, eye-tracking studies provide the weakest effects of attention on the emotional modulation of LTM, suggesting a dissociation between eye movements and neural activity. This apparent discrepancy might be explained by the diversity in stimuli used, the length of time between encoding and recall, and the experimental paradigms used to manipulate attention. Although these differences make it difficult to establish whether emotional memory benefits more from enhanced attention at encoding, or from preferential post-encoding processes, they provide testable hypotheses for future research, which we discuss below.

### 4.3. Limitations of included studies and future directions

The heterogeneity between behavioural and neuroimaging studies may be due to differences in recall intervals (immediate recall versus a 24 hour consolidation period), stimuli type (words, faces, images, and scenes) and measures of attention (DPT vs EEG), as emotional memory is sensitive to these methodological parameters (Bennion, Ford, Murray & Kensinger, 2013a; Riggs et al., 2011). Specifically, all fMRI and eye-tracking studies used emotionally valenced images, while the majority of EEG studies used emotional words. Similarly, the studies which have a recall interval of 12hrs or longer (e.g., Bennion et al., 2013b; Humphreys et al., 2010) report lower effects than studies with immediate recall. Accordingly, to determine the effect of attention on emotional LTM, stimuli type and recall intervals need to be better controlled across experiments. The field would also benefit from the direct comparison of image-based versus word-based tasks in terms of differences in electrophysiological and neuroanatomical attention-related responses at encoding, and further, determine whether memory depends on these processes, or relies more heavily on consolidation processes.

Experiments examining early sensory-related ERPs, such as the P1 and N1, reported no difference in amplitude between emotional word types. Previous research using oddball tasks containing images report significant modulations on the amplitude of the P1 and N1 components, as opposed to the passive viewing paradigms involving words typically employed when investigating attention and emotion (Carretie´ et al., 2004; Delplanque et al., 2004). As such, further research is needed that adopts attention-demanding paradigms at encoding using images rather than words. This will allow for stronger interferences to be made regarding the influence that attention-modulated, sensory-related ERPs have on emotional LTM.

It is well established that stimulus type can bias participants’ memory responses, such that emotion can modify the qualitative features of how stimuli are remembered; a change that is measureable depending on the memory variables used (e.g., familiarity, recognition accuracy; Bennion et al., 2013a). As this review did not account for specific memory performance measures, it is unclear if the behavioural and neural correlates of attention interact with qualitative changes in emotional memory. This is important to consider given that compared to familiarity-based memory, recollection involves additional prefrontal cortex activation, activation of the inferior parietal lobe and reactivation of sensory areas that are active during encoding (Skinner & Fernandes, 2007). By contrast, familiarity-based memory is associated with decreased activity of the perihinal cortex and rhinal cortex (Skinner & Fernandes, 2007), and is unaffected by reduced attention at encoding (Curran, 2004). Whether emotion interacts with familiarity- and recollection-based memory and their neural correlates under conditions of divided attention is a question needing to be addressed by future research.

Additionally, alternate EEG methods may be more suited to investigating the attention-emotion interaction. Evidence suggests oscillatory dynamics may be more reflective of attention-related cortical activity than ERPs (Yordanova, Kolev & Polich, 2001). Oscillatory activity provides useful information on the physiology of brain dynamics, with research demonstrating alpha-band activity (8 – 12 Hz) reliably indexes fluctuations in cortical excitation, and has been shown to be sensitive to emotional stimuli (Foxe & Snyder, 2011; Güntekin & Basar, 2014; Klimesch, 2012; Payne & Sekular, 2014; Uusberg et al., 2013). Studies (Aftanas et al., 2002; Güntekin & Başar, 2007; Onoda et al., 2007; Uusberg et al., 2013) have reported an increase in alpha desynchronisation in response to aversive (e.g., mutilated human bodies) compared to positive and neutral stimuli, suggesting a decrease in alpha power in response to emotional information may reflect enhanced neuronal excitation induced by affective attention, whereby attention is rendered to emotional stimuli for heightened sensory processing. This interpretation is consistent with behavioural findings of valence-induced changes in attention at encoding and superior performance for emotional stimuli at recall (MacKay et al., 2004; Riggs et al., 2011).

The adoption of alternative electrophysiological measurements, such as analyses of oscillatory activity, would be complemented by comparing attention-related memory enhancement with factors known to facilitate memory consolidation, such as sleep (Rasch & Born, 2013). Due to a disengagement from the external inputs of wakefulness, sleep is argued to provide a suitable environment for offline processes to facilitate localised synaptic downscaling and the distribution of hippocampally stored information into LTM (Ellenbogen, Hulbert, Stickgold, Dinges & Thompson-Schill, 2006; Rasch & Born, 2013). Using fMRI, Sterpenich et al. (2009) revealed sleep promoted the reorganisation of neuronal representations of emotional pictures, such that greater picture recognition accuracy after sleep compared to time awake was associated with enhanced activation of the ventral medial prefrontal cortex, an area involved in memory retrieval, and the extended amygdala, a region involved in modulating emotional information at encoding. These findings are in line with Bennion et al. (2015b), who found that that memory for emotional information was greater after a period of sleep versus wake, and this was mediated by cortisol levels at encoding. Further, the beneficial role of sleep on memory has more recently been documented with a 90 minute nap (Payne et al., 2015), suggesting that even after brief periods, sleep actively influences long-term, system-level consolidation for emotional content. However, no study has directly examined the interaction between attention and sleep on emotional memory consolidation, limiting the ability to establish whether emotional memory benefits from enhanced attention at encoding, or from post-encoding (i.e. sleep-dependent) memory consolidation.

## 5. Conclusions

This systematic review and meta-analysis aimed to determine whether attention significantly impacts memory for emotional stimuli. A secondary aim was to determine whether there was homogeneity between behavioural, electrophysiological, and neuroanatomical correlates for the effect of attention on emotional memory. Overall, attention plays a significant role in facilitating memory of emotional over neutral information, and this effect was reported across behavioural, eye-tracking and neuroimaging studies. The electrophysiological correlates of the attention-emotion memory interaction appear to be more reliably indexed by ERP components later in the processing cascade, such as the LPP; however, the role of sensory-related components may be better explored by systematically manipulating attentional demands. Neuroanatomically, increased activation of the amygdala, ventral visual stream and vmPFC were reported under conditions of divided attention, suggesting these regions interact to prioritise emotional over neutral information. However, despite extensive research, examinations of the eye-tracking and neural correlates of attention and emotional LTM are inconsistent. This inconsistency likely reflects differences in emotional stimuli, paradigms, and recall intervals, and as such, future research should better control for these methodological parameters. Further, although ERPs are useful in assessing the temporal processing of attention to emotional stimuli, future research should examine changes in oscillatory dynamics. Finally, investigating the interaction between attention and sleep may further our understanding of whether emotional memory depends on attention at encoding, enhanced consolidation processes, or an interaction between the two.

## Acknowledgements

ZRC is supported by a Research Training Program Scholarship awarded by the Australian Commonwealth Government (ID: 212190). We thank Professor Ina Bornkessel-Schlesewsky for helpful comments on an earlier version of this manuscript.

